# RSTG: Robust Generation of High Quality Spatial Transcriptomics Data using Beta Divergence Based AutoEncoder

**DOI:** 10.64898/2026.02.23.705895

**Authors:** Agrya Halder, Abhik Ghosh, Sanghamitra Bandyopadhyay

## Abstract

One of the key challenges in spatial transcriptomics data analysis is the lack of sufficient data to train models. To address this shortcoming, multiple generative models have been developed to generate synthetic spatial transcriptomics samples in a controlled environment. However, these models often fail in out-of-the-box generation in the presence of noise (such as outliers). To tackle this challenge, we propose RSTG (Robust Spatial Transcriptomic Generator), an autoencoder that incorporates the *β*-ELBO loss, to generate realistic and high-quality spatial transcriptomic sequences. Our model uncovers data’ intrinsic structure by approximating its underlying distribution through variational inference, resulting in more interpretable and robust density estimation. We validate the effectiveness of RSTG across multiple tasks, including the recovery of cellular positions in both the 2D spatial and location domains. Our method shows improved performance, both qualitatively and quantitatively, across multiple datasets, including brain and liver samples generated using MERFISH, MERSCOPE, and Visium technologies. We further illustrate the robustness of RSTG to outliers by contaminating a portion of the data with anomalies (such as white noise, batch effects, and dropouts) as well as on a reallife degraded sample. The results show that our proposal maintains high quality and stability even when the training data are contaminated, across a variety of experimental settings and in comparison with existing approaches.

## I. Introduction

THE spatial transcriptomics (ST) technology [1], [2], [3] represents a groundbreaking advancement in molecular biology. While single-cell RNA sequencing (scRNA-seq) enables high-resolution analysis of gene expression at the cellular level, it lacks information on the physical location of cells, making it difficult to understand tissue organization. ST, on the other hand, overcomes this limitation by capturing gene expression data along with the spatial positioning of cells, thus providing insights into the spatial architecture of tissues [4], [5], [6], [7], [8]. It allows researchers to gain deeper insights into how gene activity is organized across different tissue regions, how cells interact within their microenvironment, and how tissues are organized at the molecular level. Although the ST technology has advanced considerably, the acquisition of ST data remains difficult and costly. Moreover, obtaining biological samples can be challenging, and some samples may be too rare for practical analysis. Consequently, limited sample sizes may fail to capture the true characteristics of the population, potentially causing imbalances that undermine the reproducibility of experimental findings. In recent years, in silico data generation has played a significant role in computer vision, particularly in data augmentation tasks. Deep learning models such as Generative Adversarial Networks (GANs) [9], [10], [11], [12], and Variational Autoencoders (VAEs) [13],[14] have been widely employed to produce photorealistic images. GANs are composed of two competing neural networks, a generator and a discriminator, that are trained simultaneously in a game-theoretic setup, where the generator aims to produce realistic samples and the discriminator strives to distinguish between real and generated samples. The generator takes a low-dimensional distribution as input and aims to map it to high-dimensional distributions (i.e., realistic images) that are indistinguishable from the training samples. Meanwhile, the discriminator is trained to authenticate generated samples with likelihood scores. More recently, diffusion-based generative models [15] have emerged as a powerful alternative, generating high-fidelity samples through iterative denoising processes. Although these methods aim to generate highquality data, these models focus primarily on generating high-dimensional data from known distributions, failing to meet the need for supplementing unmeasured data. Additionally, these conditional generation methods often begin by clustering the dataset and then guide the generation process using cluster labels, which limits their ability to perform more detailed, fine-grained conditional generation.

While the notion of data generation has become a pivotal scope to explore, the robust generation of synthetic ST data under possibly noisy conditions is still to be investigated.

Sometimes, ST data may contain anomalies as a form of noise. When synthetic samples are generated from these noisy ST data, these anomalies can significantly alter their representation, leading to discrepancies between the generated data and the underlying patterns they aim to replicate. This can affect the quality and reliability of downstream analyses, making it challenging to derive accurate insights or build robust models. Therefore, effective noise handling is essential, and robust data generation is highly important for accurate downstream analysis of these spatial sequences. To mitigate these challenges, we introduce RSTG, a robust autoencoder on variational inference [16] to generate realistic, high quality ST sequences (see Fig. 1). Previously, multiple studies [17], [18], [19], [20] have utilized the concept of beta divergence in several learning problems to ensure robustness against different forms of contamination, facilitating consistent performance across varying conditions and datasets. However, these approaches primarily address global or unstructured noise in generic settings and do not explicitly account for the heterogeneous and structured noise characteristics inherent in ST data, such as spatially varying dropout, gene-specific sparsity, batch-induced effects, and real-world noise (such as degraded tissue). In contrast, our model not only addresses this gap by incorporating *β*-divergence–based variational inference to adaptively downweight corrupted observations, but also effectively retains downstream performance across two ST objectives, namely spatial layer recovery and 2D coordinate recovery, across datasets under varying degrees of contamination. Our model provides a computationally efficient framework compared to traditional computationally expensive diffusion-based and adversarial-based generative models while maintaining robustness to structured perturbations in ST data, including degraded tissue samples representing real-world anomalies.

**Fig. 1.**
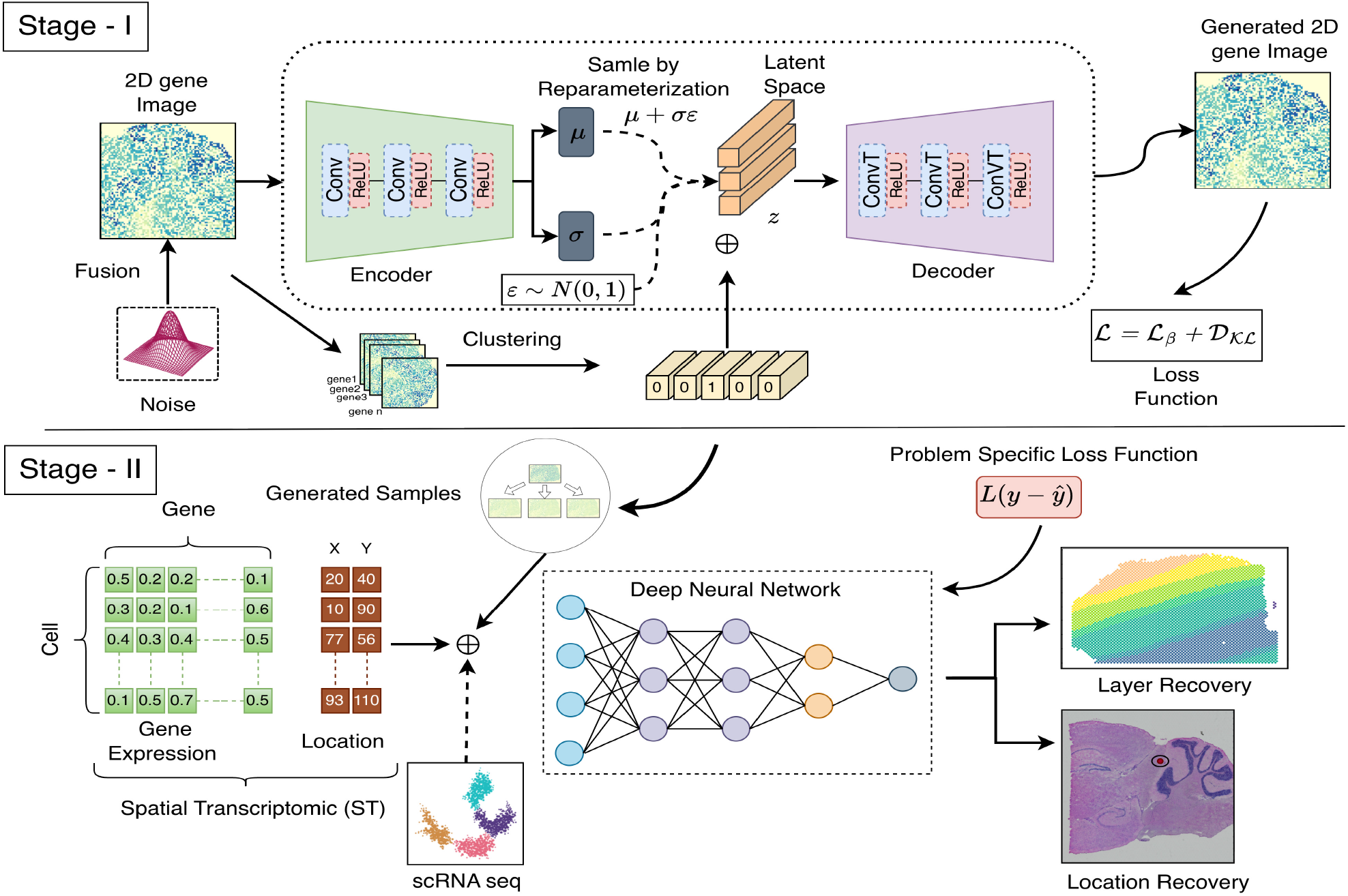
Overview of the proposed RSTG method. The proposed method consists of two stages: Stage I and Stage II. Stage I refers to the data generation approach for single-cell ST data, and Stage II represents the recovery of spatial cell locations or layer using task-specific loss functions.

The main contributions of our work are summarized as follows:

- We introduce RSTG, a VAE-based data generation approach for single-cell ST that incorporates noise-resilient strategies inspired by robust statistical theory.
- We adopt an end-to-end unified framework that jointly integrates synthetic data generation and downstream spatial analysis to capture transcriptional realism. Realistic ST sequences are first generated using a *β* divergence-based VAE that learns contamination-resilient latent representations (Stage I), and are subsequently integrated with the original observations within a single optimized pipeline to improve the spatial layer and coordinate recovery (Stage II).
- We further demonstrate that under varying forms and strength of contamination, including white Gaussian noise, dropout events, and batch effects, our model enables stable generation of ST data by successfully suppressing the adverse effects of corrupted observations.
- Through quantitative and qualitative evaluations on five public datasets, encompassing both artificial and realworld contamination, we demonstrate that RSTG surpasses existing state-of-the-art (SOTA) methods in both robust data generation and downstream analysis tasks, including spatial location and layer recovery.

## II. Literature survey

Recent advancements in generative techniques have opened up significant opportunities for in silico transcriptomic data generation [21]. The generation of spatial transcriptomic data is usually done using GANs or VAEs. scVI [22] uses a stochastic optimization technique for probabilistic representation of gene expression. Although scVI focuses on downstream analysis tasks, it does not generate comprehensive gene expression profiles of cells. The cscGAN [23], a GAN-based mdoel, was proposed to learn complex, non-linear relationships between genes across diverse, multi-cell-type datasets, enabling the generation of realistic cells belonging to specific types. However, it faces several limitations, such as excessive smoothing and a reduction in the natural variability of gene expression between individual cells. Xu et al. overcome these issues by proposing the scRNA-seq imputation GAN, scIGAN [24]. ScIGAN leverages generated cells instead of observed ones and achieves a balanced performance across both major and rare cell populations. The LSH-GAN [25], introduced by Lall et al., is a generative network that updates its training process using locality-sensitive hashing to produce new realistic cell samples.

However, these models are limited in their ability to generate samples beyond the training distribution, restricting their usefulness in supplementing unmeasured data. Furthermore, GAN-based models demand complex architectural designs, meticulous parameter tuning, and specialized optimization techniques to achieve stable training [26], [27]. These requirements create significant challenges for their application to new datasets and data generation under specific conditions, limiting their practicality and flexibility across diverse scenarios.

Recently, advancement in latent-based generative models such as VAE has demonstrated impressive performance gains in multiple domains such as video, audio, and text. Compared to GANs, VAEs offer a more stable training process for generating realistic samples from complex data distributions. However, the efficacy of these methods for single-cell data generation remains insufficiently explored.

## III. Methodology

### A. Overview

As shown in Fig. 1, in Stage I, an autoencoder is trained with a *β*-*ELBO* loss that takes as input the ST data and generates high-quality synthetic ST samples. The encoder takes the distribution of the real ST data and maps it into a reduced-dimensional latent space parametrized by *µ* and *σ*. The decoder takes the latent variable *z* (i.e., sampled from *µ* and *σ*) and maps it back to the original data space by generating a reconstructed output. A one-hot vector, representing the clustering label of genes, is concatenated with the latent space and fed through multiple covolutional layers to generate gene expression with a shape similar to the input. In Stage II, the ST samples generated by the autoencoder, together with the real samples, are used to train a DNN model for the downstream analysis. Given scRNA data with unknown cell locations, the deep learning model predicts the 2D spatial coordinates or layer information of each cell. A problem-specific loss function is used to predict either the layer or the spatial location of the given ST sample.

### B. Preprocessing: 2D Gene Embedding

In this step, the ST data are transformed by considering each gene as a separate sample. Let *G* ∈ *R*^*M×N*^ denote the gene expression matrix obtained from an ST dataset, where *M* denotes spatial locations (or spots) and *N* indicates the total number of genes. For any given gene *g*, the embedding function takes its correspondence feature vector and rearranges it into a 2D matrix *I*_*g*_ ∈ *R*^*h×w*^ based on the spatial coordinates of each spot. Here, *h* and *w* represent the number of rows and columns covering the tissue capture area, respectively. If a spot lacks a corresponding entry in the 2D gene expression matrix (missing or unmatched), its gene expression value is set to 0. The 2D embedding is formulated as,

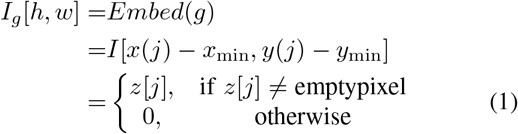

Here, *x*(*j*) and *y*[*j*] indicate the spatial coordinates of the *j*_*th*_ spot, while *z*[*j*] corresponds to the expression value of gene *i* at that location.

### C. Stage-I: Beta ELBO based Data Augmentation

Here, we construct a Variational Autoencoder framework that includes an encoder *E* and a decoder *D*, both implemented using convolutional neural networks that approximate model’s posterior distributions. Supplementary Table 1 provides the architectural details of the VAE. Specifically, the encoder transforms the input matrix *I*_*g*_ ∈ *I* into a latent representation *z* by approximating the true posterior *p*(*z I*_*g*_) with a parameterized distribution *q*_*ϕ*_(*z I*_*g*_), commonly modeled as a Gaussian distribution. The approximate posterior is defined as 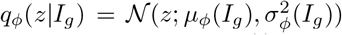, the normal density evaluated at *z*, with mean *µ*_*ϕ*_(.) and variance 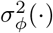 both predicted using a convolutional neural network with parameters *ϕ*. The decoder transforms the latent representation *z* back into the original data space by approximating the conditional distribution *p*_*θ*_(*I*_*g*_ *z*). This reconstruction process is modeled using a neural network parameterized by *θ* to recover the input *I*_*g*_ from the latent code. The output from the decoder D is assumed to follow a Gaussian distribution to capture reconstruction uncertainty. The overall marginal likelihood of the observed data under this generative model is expressed as:

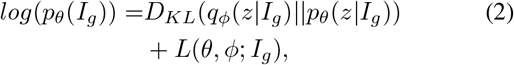

where, *log*(*p*_*θ*_(*I*_*g*_)) denotes the log-likelihood of the observed input *I*_*g*_, *D*_*KL*_(*q*_*ϕ*_(*z I*_*g*_) ∥*p*_*θ*_(*z I*_*g*_)) represents the Kull(*p* ), defined as and the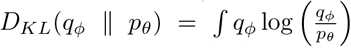 back–Leibler (KL) divergence, which measures the divergence between the approximate posterior (*q*_*ϕ*_) and the true posterior second component *L*(*θ, ϕ*; *I*_*g*_), called the Evidence Lower Bound, serves as a surrogate objective for training. The ELBO is further defined as:

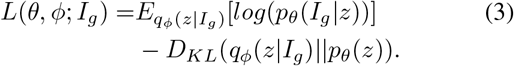

The first term in (3) quantifies the expected log-likelihood and acts as the reconstruction loss, while the second term (KL divergence) serves as a regularization term that encourages the approximate posterior to remain close to the prior distribution. We now introduce a robust formulation of variational inference, inspired by the principles outlined in [16], to enhance the resilience of variational autoencoders to outliers. In this framework, we define a *β*-cross entropy 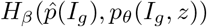, which measures how much the model’s generative distribution *p*_*θ*_(*I*_*g*_ *z*) deviates from the empirical data distribution 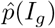 for each input *I*_*g*_. The *β*-ELBO, considering the regularization assumption on latent variables as standard Gaussian𝒩 (0, 1), is defined as:

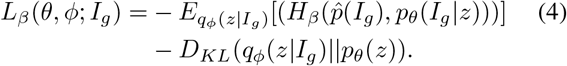

In the Gaussian setting, the likelihood *p*(*I*_*g*_ | *z*) is modelled as a normal distribution 𝒩 (*Î*_*g*_, *σ*), where *Î*_*g*_ denotes the decoder’s output reconstructed from the latent variable *z* and *β*-cross entropy is defined as:

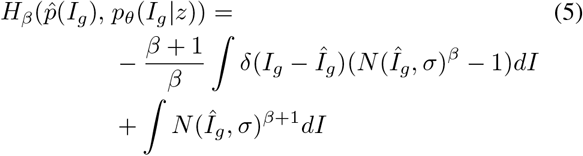

In Eq. (5), the second term is independent of *Î*_*g*_, and therefore does not influence the optimization with respect to the decoder’s output. Consequently, minimizing the *β*-cross entropy reduces to maximizing the term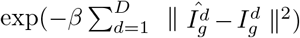, where *D* denotes the dimensionality of the input. Therefore, the final simplified loss function can be expressed as:

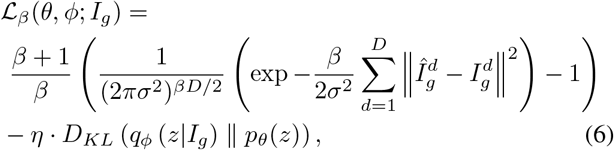

The first term in (6) corresponds to the *β*-cross entropy loss, incorporating a robust exponential measure of discrepancy between the reconstructed output *Î*_*g*_ and the original input *I*_*g*_. The second term is the KL divergence term, which again acts as a regularizer. The loss function is scaled by a robustness hyperparameter *β*, which controls the influence of outliers during generation. Unlike standard ELBO, which employs a quadratic penalty MSE, the exponential term in Eq. (6) provides a heavy-tail robustness against outliers. In the ST context, when a gene matrix *I*_*g*_ contains anomalous noise (or artifacts), the reconstruction error ∥*Î*_*g*_ −*I*_*g*_ ∥^2^ becomes large.

When *β >* 0, the exponential term rapidly decays toward zero, effectively reducing the contribution of that outlier to the total gradient. This preserves the underlying biological characteristics shared across the tissue while suppressing localized artifacts. In our experiments, we set *σ* = 0.5, as the input values were normalized to lie within the range [0, 1], making this suitable for the Gaussian reconstruction assumption. The parameter *η* serves as a weighting factor to balance the contribution of the reconstruction loss and the KL divergence term. Unless otherwise specified, *η* is set to 1 to maintain comparable scales between the two loss components and ensure stable optimization.

The augmentation framework consists of three key stages. First, each gene’s expression profile is reshaped from a 1D vector into a 2D spatial matrix based on the spot coordinates, ensuring alignment with the spatial layout of the tissue (during pre-processing). Second, K-means clustering is performed on these spatial gene matrices to group genes exhibiting similar expression patterns. The cluster assignments are encoded as one-hot vectors, and the optimal number of clusters *K* is determined empirically using a rate-distortion plot (shown in Supplementary file, Fig. 1), evaluating the trade-off between cluster compactness and model complexity. Third, a variational autoencoder is trained to learn a compressed representation of each gene. The encoder outputs mean *µ* and variance *σ*^2^, which are used to sample latent embeddings from the distribution 𝒩 (*µ, σ*^2^). These embeddings, concatenated with the one-hot cluster vectors, are decoded to reconstruct the spatial gene expression matrices. The synthetic reconstructions are then combined with the original ST data to expand the training set for downstream analysis. Detailed visualization of synthetic gene samples is given in the supplementary Fig. 2. The overall augmentation procedure is described in the Supplementary File, Algorithm I.

### D. Stage-II: Spatial Location Recovery

The spatial location recovery involves predicting the spatial coordinates or layer assignment of a spot from its gene expression profile.

#### 1) 2D coordinate recovery

Let *S*_*i*_ represents the gene expression of the *i*^*th*^ spot associated with true coordinates (*x*_*i*_, *y*_*i*_). The spatial coordinate recovery model takes the *i*^*th*^ gene expression and its corresponding coordinates as input and trains a Deep Neural Network (DNN). This network captures the underlying spatial relationships present within the ST dataset. The trained model then predicts unknown coordinates from the gene expression profile of a spot. The DNN architecture begins with an input layer aligned with the dimensionality of the input gene expression vector *S*, followed by several hidden layers that progressively reduce in size, thereby enabling the extraction of hierarchical features. The output layer has dimensionality 2, representing the predicted spatial coordinates 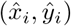 for each spot. The parameters of the DNN model are optimized with an Adam optimizer to minimize the following Mean Squared Error (MSE) loss:

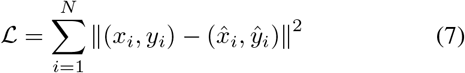

#### 2) Spatial Layer Recovery

To predict the spatial domain/layer assignment, the supervised DNN takes as input the gene expression of the *i*^*th*^ spot along with the domain label *d*_*l*_∈ *L*; *l* = 1, 2, …, *L*. The model learns the relationship between domain labels and gene expression using a logistic loss function similar to that in [28]. The logistic loss is defined as follows:

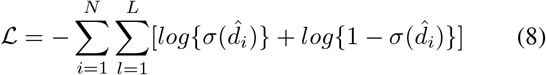

where 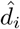 is the predicted domain for the *i*^*th*^ gene expression profile, 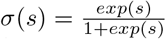, and *L* is the total number of distinct spatial domains.

For ordered domain recovery, such as predicting cortical layers in brain tissue, we adopt a rank-consistent logistic regression approach that captures ordinal relationships between domain labels. The corresponding loss is defined as,

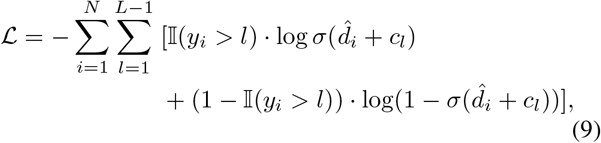

where *c*_*l*_ is a learned threshold, separating domain *l* from domain *l* + 1 with *y*_*i*_ being the layer index.

## IV. Experiments

### A. Datasets and Pre-processing

We consider a diverse collection of five public datasets representing heterogeneous tissue types: (1) Human Dorsolateral Prefrontal Cortex (DLPFC) [29], (2) Mouse posterior brain [30], (3) Mouse brain MERFISH [31], (4) Xenium breast cancer [32], and (5) Human liver cancer MERSCOPE [33]. These datasets are selected to represent a broad spectrum of biological and technical variability, spanning different tissues, species, and ST platforms. The LIBD DLPFC and Xenium datasets exhibit structured cortical and tumor microenvironments, respectively, while the MERFISH dataset offers singlecell resolution of neuronal heterogeneity. The human liver cancer MERSCOPE dataset is an FFPE-derived ST dataset, that captures real-world tissue degradation. For pre-processing, we follow a similar normalization strategy across all datasets, where we first perform spot-level normalization independently for each sample, followed by a logarithmic transformation to stabilize variance. A detailed overview of the experimental datasets is available in the Supplementary file, Table I, and Content 1.

**Table I.**
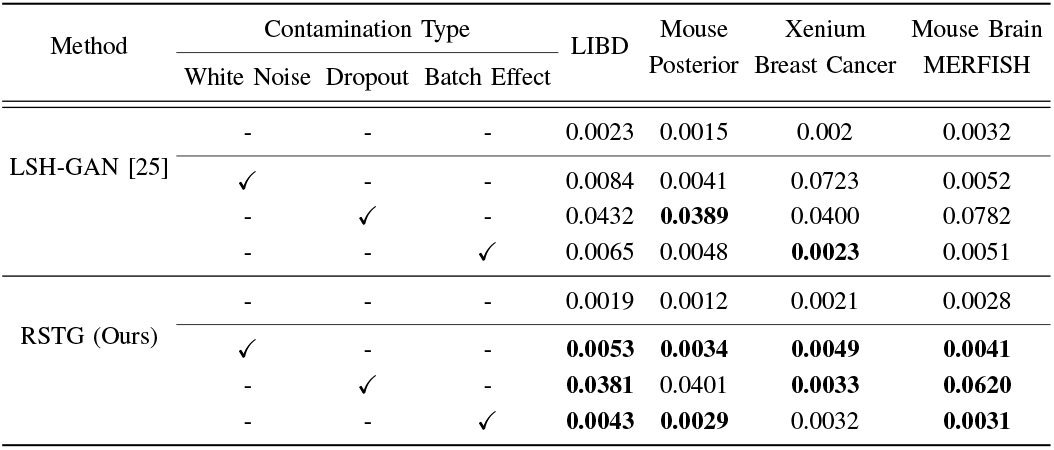
Wasserstein distance between generated and real samples across contamination types on four benchmark datasets at (5%) contamination. Lower values indicate better performance.

### B. Evaluation Strategy

While evaluating single-cell generation is difficult, we use three commonly used metrics: UMAP, Wasserstein distance, and classification performance scores. These metrics serve as both quantitative and qualitative measures for the quality of synthesized cells [25], [28]. To assess downstream task performance, we consider two objectives in ST analysis: spatial layer recovery and coordinate recovery. We use Top1 and Top-2 accuracy for the LIBD dataset, which reflects the correctness of spatial layer assignments. Top-1 accuracy measures the proportion of spots correctly assigned to their true layer, while Top-2 accuracy allows for predictions that match either the true or adjacent layer. For other datasets, we report Pearson correlation, which assesses the linear agreement between the predicted and actual layer assignments. These evaluation metrics are driven by the availability and nature of dataset-specific ground truth rather than methodological inconsistency.

### C. Implementation Details

For the LIBD dataset, we evaluate our model under multiple experimental scenarios to assess both within-sample consistency and cross-sample generalization. Specifically, we consider three settings: (1) Within-Brain (scenario S1), where the model is trained on sample ID 151673 and tested on sample ID 151676 from the same brain; (2) Cross-Brain (scenario S2), where the training data remain the same (151673) but testing is performed on sample ID 151507 from a different brain; and (3) Random-Source Random-Target (scenario S3), where the model is trained on one section and evaluated on randomly selected sections from different brains (e.g., Trained on 151510, and tested on 151670). For the Mouse Posterior Brain dataset, we randomly partition the data into a 70 : 30 split, where 70% is used for training, and the remaining 30%is reserved for testing. The Mouse Brain MERFISH data are partitioned based on anatomical hemispheres: one hemisphere is used for training, while the other serves as the test set, and the process is repeated in reverse for cross-validation. For the Xenium breast cancer dataset, we use a 90 : 10 split for training and testing. For the Human liver cancer, data points exhibiting characteristic degradation markers (exceptionally low total transcript counts and high dropout rates) are flagged as degraded samples and used for training.

The proposed model is trained using the Adam optimizer for all datasets with 150 epochs, including 20 initial warmup epochs. The default batch size is set to 64 for training the augmentation model and 128 for the downstream analysis model. A fixed learning rate of 0.001 is used throughout. All experiments are performed on a single NVIDIA RTX A5000 GPU with 24 GB of memory. In our experiments, we select *β* values of 0.005, 0.01, and 0.03 to evaluate the robustness performance in our augmentation framework. These values are chosen to represent a range of low divergence adjustments, allowing us to control the influence of outliers during training while preserving the essential structure of the data. A smaller *β* (e.g., 0.005) ensures minimal deviation from the standard VAE behavior. Slightly higher values, like 0.01 and 0.03 introduce stronger resistance to outlier influence by reducing their contribution to the variational objective. To evaluate the model’s behavior under diverse contamination scenarios, we implement three distinct types of synthetic noise that are commonly encountered in practice. White noise contamination is introduced by injecting random Gaussian noise into a subset of samples to simulate measurement errors. Batch effects are simulated by adding gene-specific biases uniformly across all samples, mimicking systematic shifts between experimental conditions. Dropout noise is applied by randomly zeroing out a proportion of gene expression values, reflecting missing data common in single-cell experiments. This experimental setup allowed us to assess the model’s capacity to generalize across varying noise conditions. The selected *β* values were empirically validated to ensure model stability and consistent performance across datasets, demonstrating the framework’s robustness without requiring explicit supervision or fine-tuning.

### D. Baseline

We categorize the baseline methods into two groups: augmentation-based approaches and spatial recovery approaches. The augmentation group includes several benchmarks methods, such as LSH-GAN [25], cscGAN, splatter, SUGAR, traditional GAN, f-GAN, and W-GAN. Among the augmentation methods, LSH-GAN has demonstrated superior performance over other generative models (refer to [25]), and thus we consider it as our primary competitor for comparing ST data generation. For spatial location recovery task, we consider Tangram [34], SpaOTsc [35], novoSpaRc [36], and CeLEry [28] as representative baselines.

## V. Results and Discussion

### A. Comparison with State-of-the-Art Methods

#### 1) Generation Quality

The results presented in Table I show a quantitative comparison between RSTG and LSH-GAN on four public datasets with varying contamination types. From these, it is evident that RSTG consistently outperforms LSH-GAN in terms of Wasserstein distance between generated and real samples. Under white noise contamination in the Breast Cancer dataset, RSTG demonstrates strong robustness against additive noise, where the Wasserstein distance drops significantly from 0.0723 in LSH-GAN to 0.0049. In the case of dropout, which simulates missing data in ST, RSTG further outperforms LSH-GAN in most of the datasets, with a noticeable reduction to 0.0033 in the Xenium dataset, from 0.040 of LSH-GAN. The same trend is visible in three out of four datasets, under batch-effect contamination reflecting systemic bias. However, LSH-GAN slightly outperforms RSTG for the Mouse Posterior dataset under dropout contamination, likely due to the better preservation of the sparse local expression patterns of GAN.

Overall, the results shown in Table I highlight RSTG’s superior generalization and resilience across a range of biologically realistic noise settings, making it more effective than LSH-GAN in generating high-fidelity synthetic ST samples for downstream analysis [37]. To further evaluate the quality of the generated data, we have also employed UMAP visualization for analysis, presented in Supplementary file, Fig. 3. These UMAP visualizations further demonstrate that the proposed RSTG method more effectively preserves the spatial and structural integrity of the original data compared to LSHGAN. Although the original datasets exhibit well-separated and biologically meaningful clusters, RSTG closely replicates these patterns, with clear boundaries and low noise, compared to often overlapping clusters and distorted spatial organization observed in LSH-GAN, reflecting higher fidelity of RSTG in reconstructing complex ST structures.

#### 2) Predictors Performance on Generated Samples

We evaluate the performance of generated samples from RSTG using a unified evaluation pipeline that combines generative models with a consistent set of downstream predictors, enabling evaluation of both generative quality and downstream utility. For example, synthetic ST data generated by LSH-GAN are concatenated with the original observations in a controlled ratio (1 : *n*, where *n* ∈ 2, 3, 4) to train the CeLEry ST location predictor. As shown in Table VI, the ratio of real and synthetic samples is empirically selected to explore the best trade-off between robustness and performance without introducing excessive distributional shifts (see Section V-C).

#### Spatial Layer Recovery

Regarding cortical layer recovery on the LIBD dataset, as presented in Table II (Scenario S1), RSTG achieves Top-1 and Top-2 accuracies of 66.4% and 93.5%, respectively, outperforming the strongest baseline, CeLEry, by 12.6% and 4.3%. Under 5% white noise contamination, RSTG further maintains strong robustness with Top-1 and Top-2 accuracies of 33.5% and 82.7%, respectively. Even at 10% contamination, our method preserves a high Top-2 accuracy of 75.4%, demonstrating resilience against increasing noise levels. Similarly, in Scenario S2 and Scenario S3, RSTG consistently outperforms competing approaches in both Top-1 and Top-2 accuracy across all contamination levels. For example, in Scenario S3, RSTG maintains the highest performance across all contamination settings, reaching Top-1 and Top-2 accuracies of 41.7% and 73.1%, respectively, under 10% contamination. From a clinical standpoint, these results shape the potential of RSTG to serve as a reliable data augmentation framework in spatially resolved single-cell studies by enabling accurate layer and region identification even in the presence of biological artifacts. Supplementary

**TABLE II.**
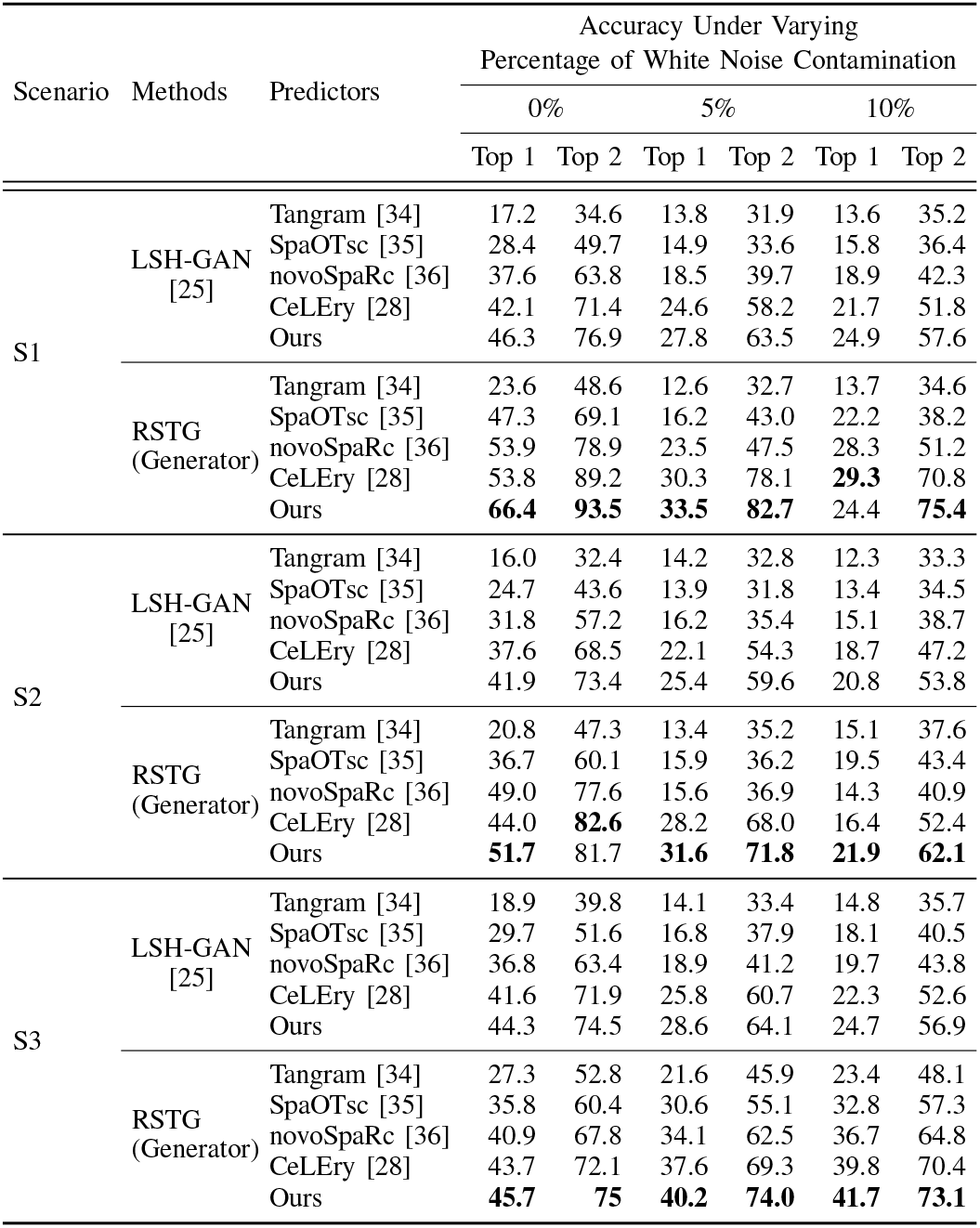
Top-1 and Top-2 classification accuracy (%) on the libd dataset under increasing white noise contamination levels (0%, 5%, 10%). Higher values indicate better performance. ***β*** is taken as **0.005**.

Fig. 4 further shows the predicted cortical layers and accuracy metrics for LIBD dataset.

#### Spatial Coordinate Recovery

Similarly, in the spatial coordinate recovery analysis, presented in Table III, RSTG continues to outperform other methods, obtaining correlation scores of 0.925 and 0.732 on the mouse posterior and MER-FISH datasets under noise-free conditions, respectively. As noise increases, the performance of other predictors such as Tangram, SpaOTsc, and novoSParc deteriorates substantially, with correlations dropping as low as 0.175 and 0.04 in mouse posterior and mouse brain merfish datasets, respectively. In contrast, RSTG demonstrates robustness to outliers, achieving consistent performance gains, while maintaining the highest values of 0.993 and 0.732 in both datasets. Even with 10% outliers, where other methods suffer heavily, our method achieves consistently higher correlations exceeding 0.974 on the mouse posterior dataset and 0.728 on MERFISH. These results highlight the superior performance and robustness of RSTG under challenging contamination levels. Supplementary Fig. 5 and 6 visualize the reconstructed gene maps of generated samples from all baselines. The computational load, represented by the number of Floating-point Operations per Second (FLOPS), further demonstrates improved spatial reconstruction performance while maintaining a competitive computational burden in comparison to SOTA methods across datasets (Supplementary file, Table IV).

**TABLE III.**
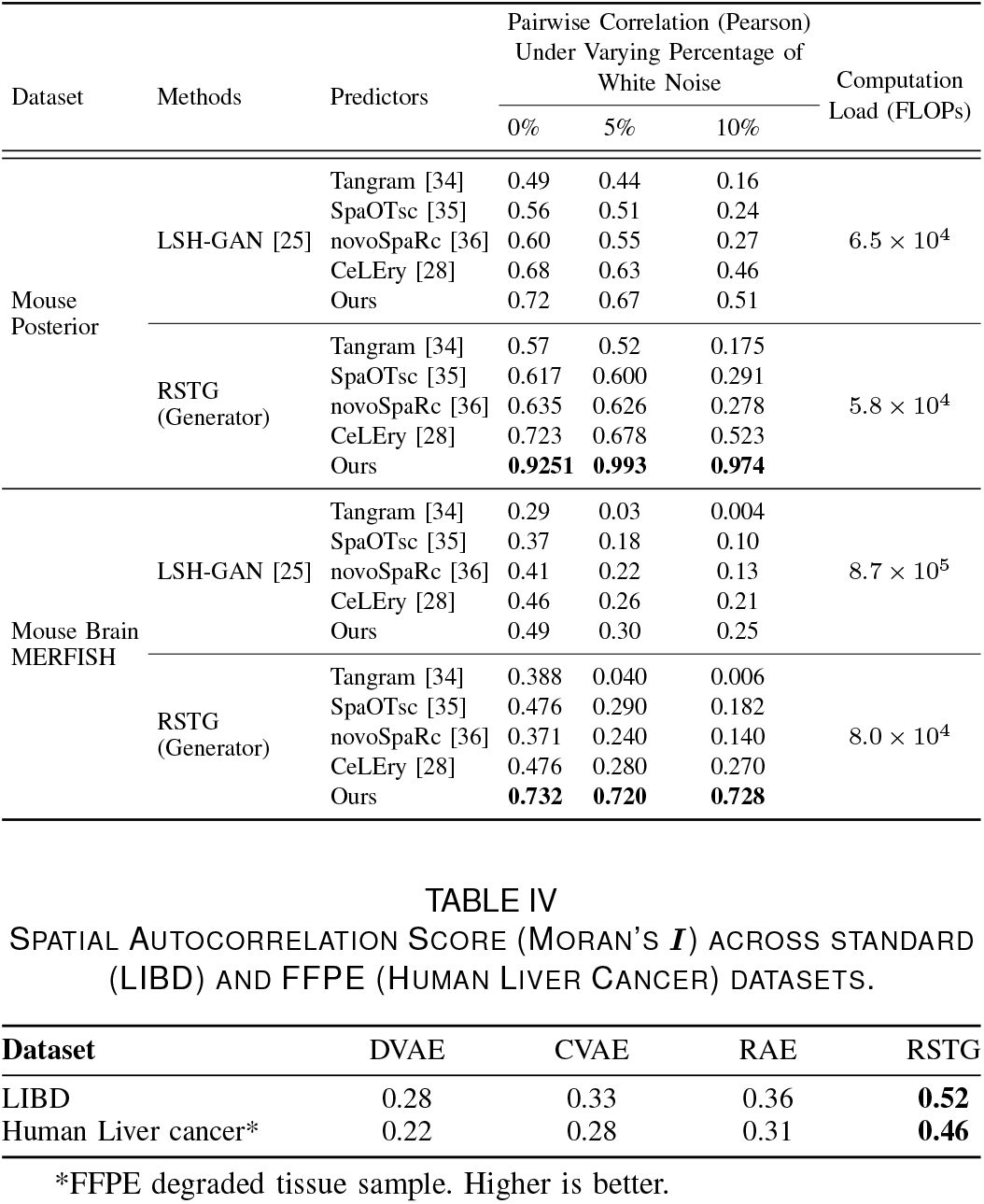
Pairwise (pearson) correlation between predicted and true coordinates on Mouse Posterior and Mouse MERFISH datasets under varying contamination levels (0%, 5%, 10%). Higher values indicate better reconstruction (***β***=0.005).

**TABLE IV.**
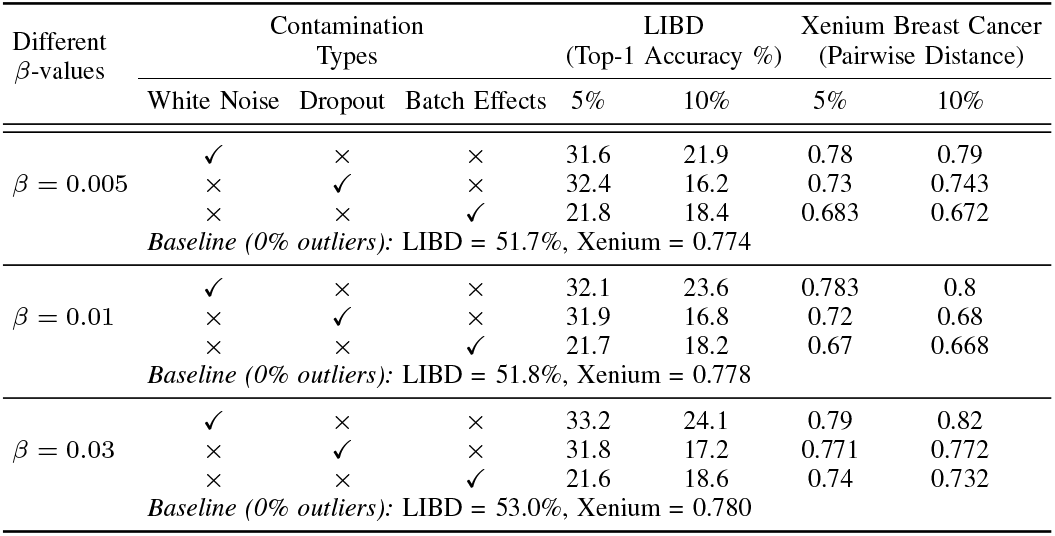
Spatial Autocorrelation Score (MORAN’S ***I*** ) across standard (LIBD) and FFPE (Human Liver Cancer) datasets.

#### 3) Comparison with Other Methods

We compare the biological validity of the samples generated by RSTG with Denoising VAE (DVAE) [38], Robust AE (RAE) [39], and Coupled-VAE (CVAE) [40]. Then, we evaluate them using spatial autocorrelation (Moran’s *I*), as a proxy for preservation of spatial biological structure. We use two different datasets: LIBD and Human Liver Cancer, where the Human Liver Cancer setting explicitly simulates realistic contamination with FFPE-like degradation by selecting low-count and high-dropout cells and LIBD simulates artificial contamination with 10% batch effects. As shown in Table IV, DVAE, which corresponds to a standard denoising VAE baseline, achieves the lowest Moran’s *I* values (0.28 on LIBD and 0.22 on Human Liver Cancer), reflecting limited capability in preserving spatial dependencies under heavy corruption. CVAE improves upon DVAE through its conditional latent structure, while RAE further enhances robustness by explicitly stabilizing reconstruction under perturbations. However, RSTG achieves the highest spatial autocorrelation (0.52 on LIBD and 0.46 on Human Liver Cancer), demonstrating superior recovery of spatial tissue coherence even under severe FFPE-like degradation. Table III in Supplementary file further reports the performance evaluation, along with MSE loss.

### B. Ablation Study on Effects of Beta values

To systematically assess the effect of *β*-values on preserving biological fidelity under contamination, we performed an ablation study under three types of contamination, namely white noise, dropout, and batch effects at varying levels of severity (0%, 5%, and 10%). The results, shown in Table V, reveal that the proposed method performs remarkably consistently across all *β* values on both datasets. For instance, at *β* = 0.005, the LIBD Top-1 accuracy remains relatively stable across contamination levels, decreasing from 51.7% at 0% contamination to 21.9% at 10% contamination. Moreover, the pairwise correlation in the Xenium dataset increases significantly, reaching up to 0.79 at 10% white noise, suggesting improved spatial coherence in the generated data. Among all evaluated *β* configurations, *β*=0.03 yields the most consistent and superior results. In the LIBD dataset, it achieves the highest Top-1 accuracy under Dropout (54.2%) and maintains strong performance across all contamination scenarios. The Xenium dataset also shows substantial improvement, with the pairwise distance correlation peaking at 0.82 under 10% contamination, indicating the highest degree of structural fidelity. Collectively, these results highlight the efficacy of *β*-divergence in enhancing model robustness against various forms of contamination. As *β* increases, the model exhibits greater resilience, particularly under Dropout and Batch Effects. Although *β*=0.03 emerges as the most optimal configuration, consistently outperforming lower *β* values, tuning *β* improves the robustness of the latent representation by reducing the sensitivity of the model to corrupted observations.

**Table V.**
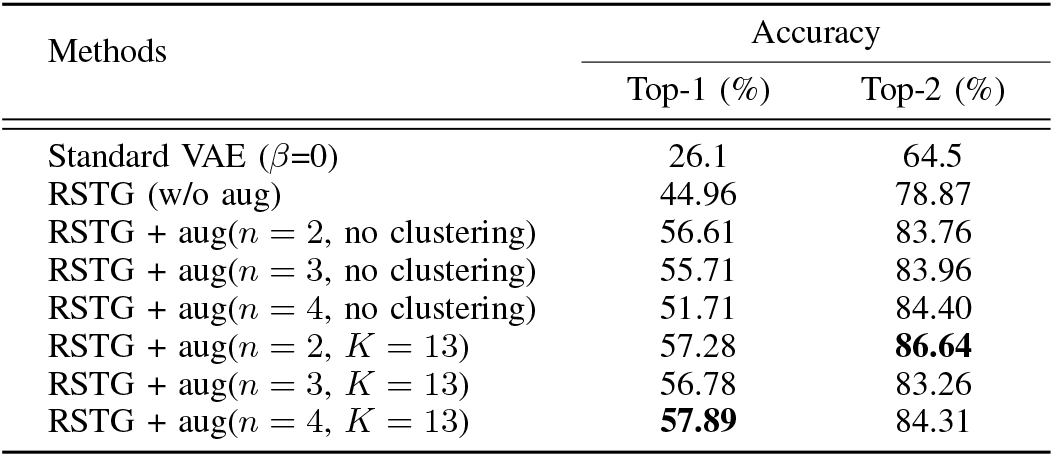
Ablation study on effects of ***β*** values with various contamination types and levels on Spatial Layer and coordinate recovery tasks on two datasets.

### C. Influence of Clustering and Augmentation in Layer Recovery

We investigate the influence of (i) the proposed *β*divergence objective, (ii) cluster-conditioned decoding, and (iii) No. of synthetic sample augmentation (n) on the spatial layer recovery task by varying the number of synthetic samples in both cases, with clustering (*K* = 13) and without clustering cases, for the layer recovery task. As shown in Table VI, the standard VAE (*β* = 0) without clustering achieves substantially lower performance (Top-1: 26.1%, Top-2: 64.5%) compared to RSTG framework without augmentation (Top-1: 44.96%, Top-2: 78.87%), demonstrating that the *β*-divergence formulation itself contributes strongly to robust representation learning and noise-tolerant reconstruction. Nevertheless, since *n* represents the number of synthetic observations generated by RSTG, choosing an appropriate value remains important for achieving optimal performance. Under the augmentation factor *n* = 2, the clustered variant (*K* = 13) improves the Top-2 accuracy from 83.76% to 86.64%. Importantly, the improvements are gradual rather than trivial, with moderate augmentation consistently improving performance compared to the non-augmented setting, indicating that clustering acts as a regularization mechanism and that synthetic data improve robustness under noisy conditions. However, performance gains do not increase monotonically with larger augmentation ratios, suggesting that overly large synthetic proportions may introduce distributional bias.

**TABLE VI.**
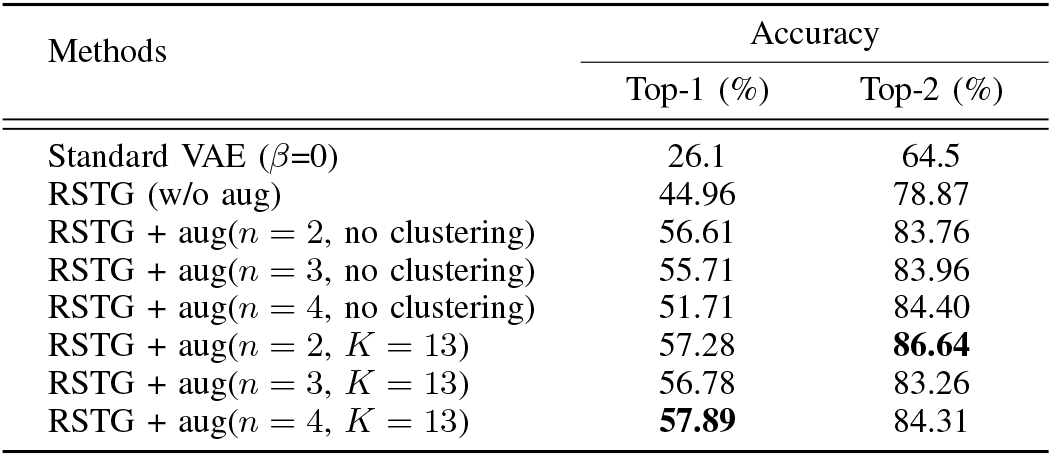
Influence of cluster-conditioned and augmentation in layer recovery performance on the LIBD dataset under 5% white noise contamination. Higher values indicate better performance.

## VI. Conclusion

This paper introduces RSTG, a robust spatial transcriptomics generation framework that integrates autoencoding and variational inference grounded in robust statistics. RSTG is designed to generate high-fidelity spatial transcriptomics data under various noise conditions. By accepting contaminated inputs that simulate anomalous or noisy measurements, the model effectively mitigates distortion and reconstructs clearer, biologically meaningful representations. The augmentation component enables the expansion of training data, improving performance in downstream analysis tasks such as spatial layer and 2D location recovery, particularly in small samples. In the second stage, a deep neural network is trained on the generated samples to enhance recovery of spatial coordinates and domain labels. Through extensive benchmarking across diverse datasets and contamination types, including white noise, dropout, and batch effects, RSTG consistently outperforms SOTA methods, such as LSH-GAN, CeLEry, Tangram, spaOTsc, and novoSpaRcin, in both data generation and spatial localization recovery tasks. While RSTG shows strong performance, it does not explicitly model rare cell types as part of the noise. It is important to note that RSTG is potentially extendable to other spatial omics modalities, such as spatial DNA methylation and RNA editing, by introducing modality-specific likelihoods, while retaining the same inference framework. Future work can address these gaps to enhance generalization further and validate RSTG across diverse spatial omics settings.

## Acknowledgment

The research work was supported by J.C. Bose Fellowship Grant JBR/2021/000036 from DST, Govt. of India. AH and SB gratefully acknowledge the support received under this fellowship during the course of this work.

