## Supplementary File for "RSTG: Robust Generation of High Quality Spatial Transcriptomics Data using Beta Divergence Based AutoEncoder"

**Supplementary Algorithm I. RSTG: Robust Data Augmentation**

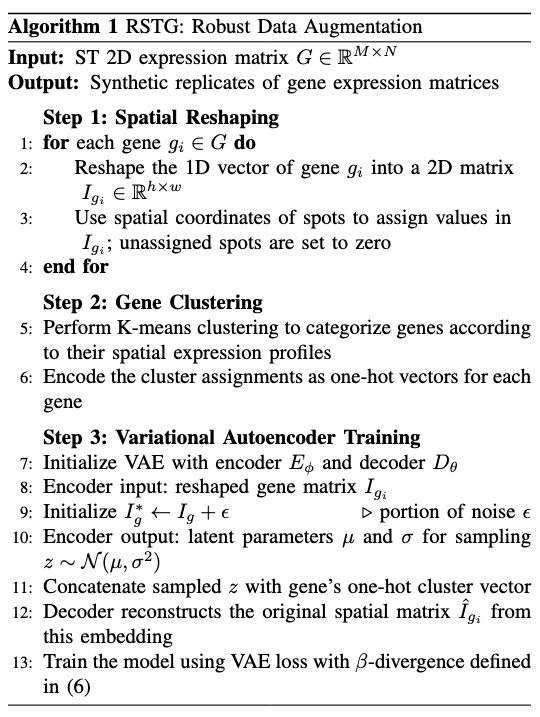

**Supplementary Table I.** Architectural View of the Autoencoder

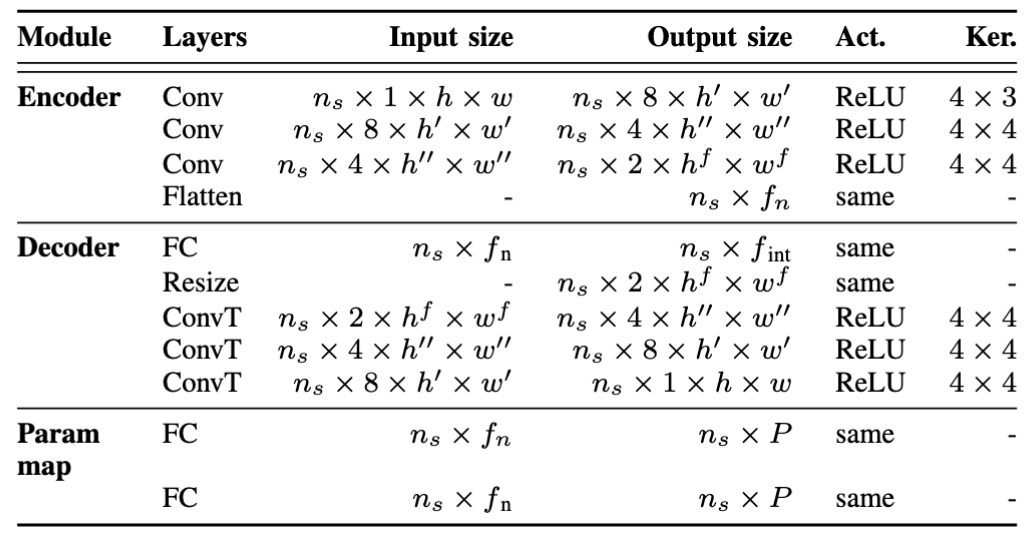

**Supplementary Table II.** A Brief overview of experimental datasets

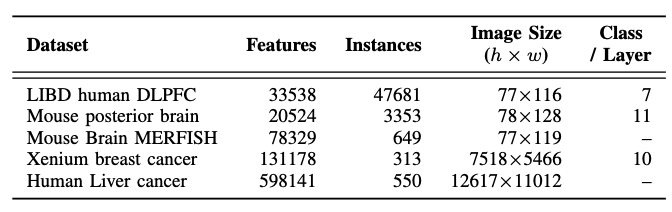

**Supplementary Table III.** Ablation study on effects of MSE with various contamination types and levels on Spatial Layer and coordinate recovery tasks on two datasets.

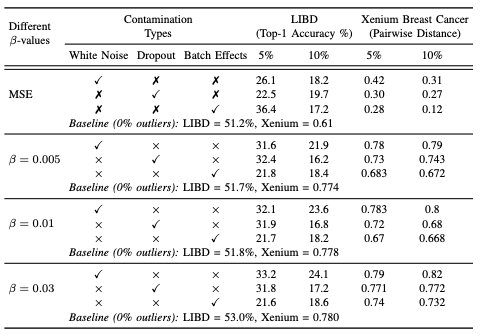

**Supplementary Table IV.** Performance comparison of different methods across datasets. Training and Inference time is reported as mean +- std (in seconds). Lower values indicate less computationally expensive.

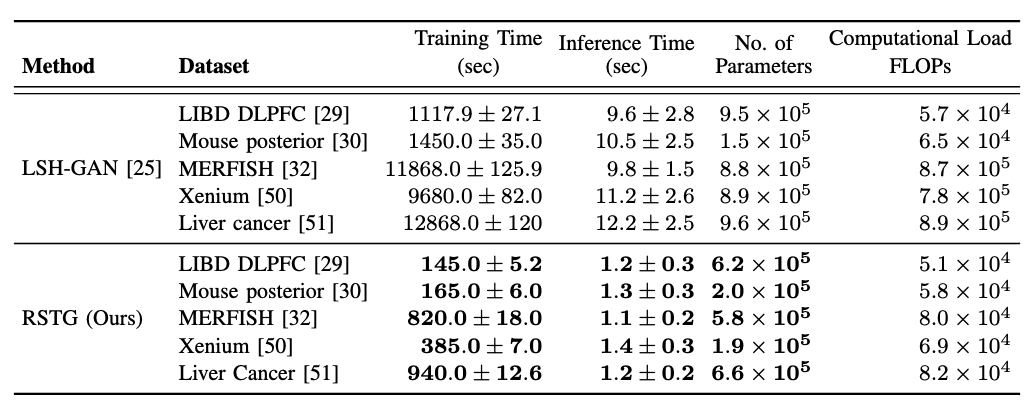

**Supplementary Content 1. Experiments: Datasets and Pre-procesing**

LIBD Human Dorsolateral Prefrontal Cortex(DLPFC) Dataset: We used publicly available LIBD DLPFC [29] ST dataset generated with 10x Genomics Visium Platform from three postmortem human brains. The dataset contains 12 multiple tissue sections, containing 33538 cells and 47681 genes, with manually annotated cortical layers (L1–L6 and WM) as raw UMI count matrices organized as spot-by-gene expression profiles along with corresponding spatial coordinates. For preprocessing, we first performed spot-level normalization to independent sample, followed by a logarithmic transformation to stabilize variance. Subsequently, we retain genes shared across all sections and merge the expression matrices together with their layer annotations to construct a unified dataset for training and evaluating scenario.

Mouse Posterior Brain Dataset: We used the mouse posterior brain ST dataset [30] generated using the 10x Genomics Visium platform. Compared to the LIBD dataset, mouse posterior brain data exhibit relatively higher spatial resolution and more detailed spatial structure with continuous spatial variation, making the reconstruction and generation task more challenging. A total 205224 spots and 3353 genes with the corresponding 2D spatial coordinates for each spot is captured here. To further enhance spatial resolution, we adopted a super-resolution strategy using TESLA [31], where gene expression is inferred at a finer grid of super-pixels (50 × 50 resolution). For downstream analysis, we identified a subset of 358 spatially variable genes using SpaGCN.

Mouse Brain MERFISH data: We used another high resolution Mouse3 sagittal ST dataset introduced by Zhang et al. [32]. In total, MERFISH imaging was performed on 1124–1147 genes in 245 brain slices (217 coronal and 28 sagittal) from four animals, generating raw molecule counts per gene per cell with associated 2D spatial coordinates. The dimension (spot-gene) of the data set is 78329 × 649. For preprocessing, we follow a similar normalization strategy as in other datasets.

Xenium Breast Cancer dataset: We used a human breast tumor ST dataset generated using 10x Xenium, which corresponds to a HER2-positive, ESR1-positive, and PGR-negative breast cancer sample in two consecutively sectioned tissue slices from the same patient. Both tissue sections (Replicate 1 and Replicate 2) measure a shared panel of 313genes with precise spatial coordinates. We particularly used Replicate 2 contains 131,178 cells for training and evaluation purpose due to better tumor heterogeneity.

Human Liver Cancer dataset: We further used a human liver cancer ST dataset generated using the MERSCOPE. The dataset is taken from FFPE tissue sections, which is known for its technical variability due to RNA degradation and fixation artifacts, making this dataset a challenging and practical benchmark. The dataset provides single-cell–resolution gene expression profiles with spatial coordinates, consisting of 598,141 cells and 550 genes. To better understand the real-world degradation, we performed basic quality control by filtering cells with very low counts (min = 10) and genes expressed in fewer than five cells, which retains the degraded and sparse nature of the data.

**Supplementary Figure 1.** Overview of rate distortion curve to find the optimal k value in clustering method. Sample ID 151507 from LIBD dataset is taken as training.

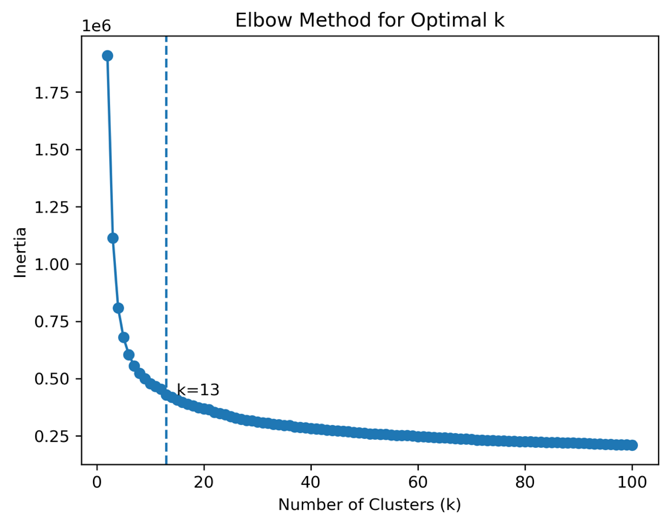

**Supplementary Figure 2.** Overview of generated samples under white noise contamination (5% and 10%). [a,d,g] show the original and generated samples without outliers. [b,e,h] correspond to 5% contamination, while [c,f,i] correspond to 10% contamination, illustrating the impact on autoencoder reconstructions.

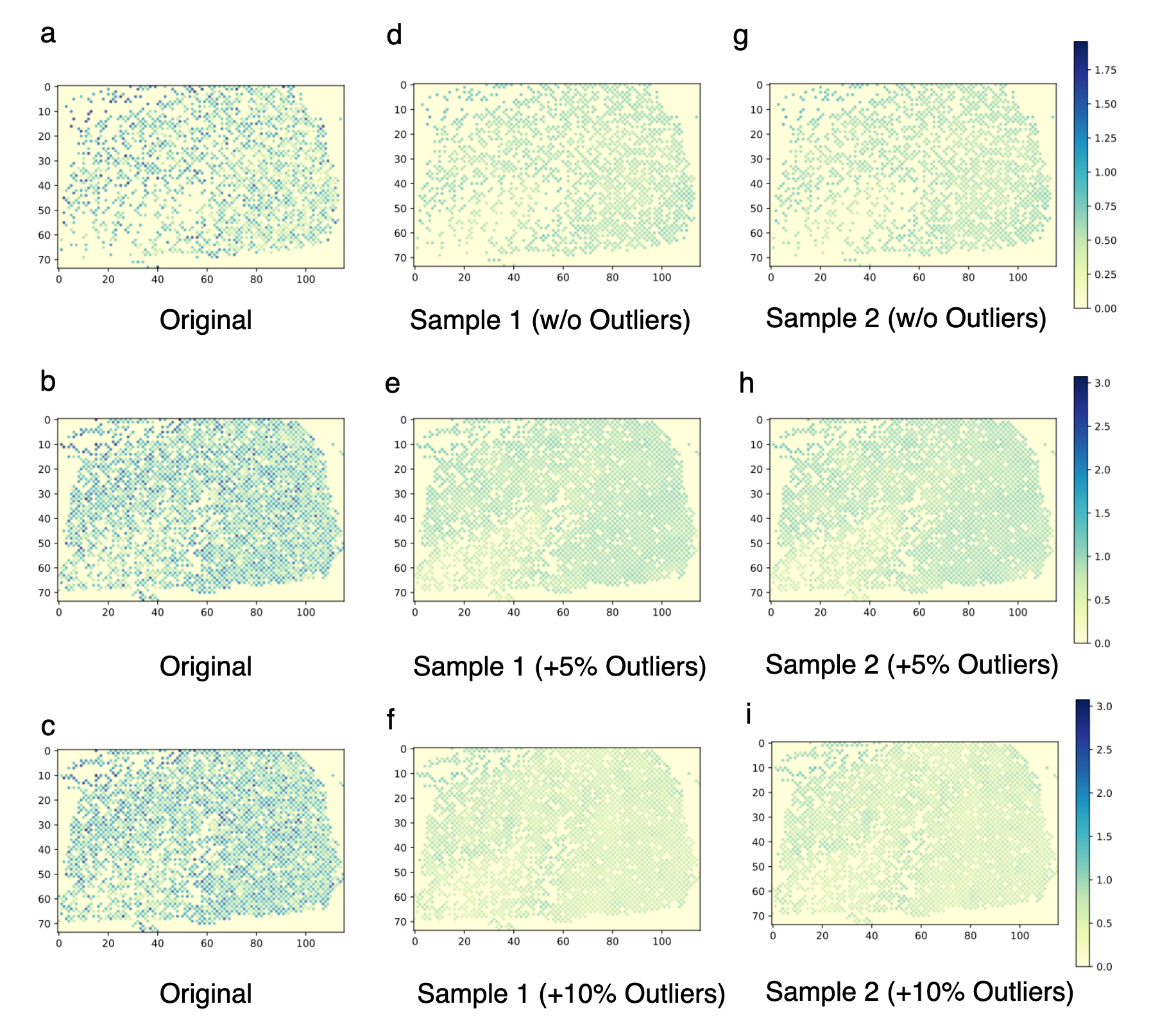

**Supplementary Figure 3.** UMAP plots illustrating the spatial distribution of cells in the original datasets, and those reconstructed by LSH-GAN and the proposed RSTG method across four benchmark datasets.

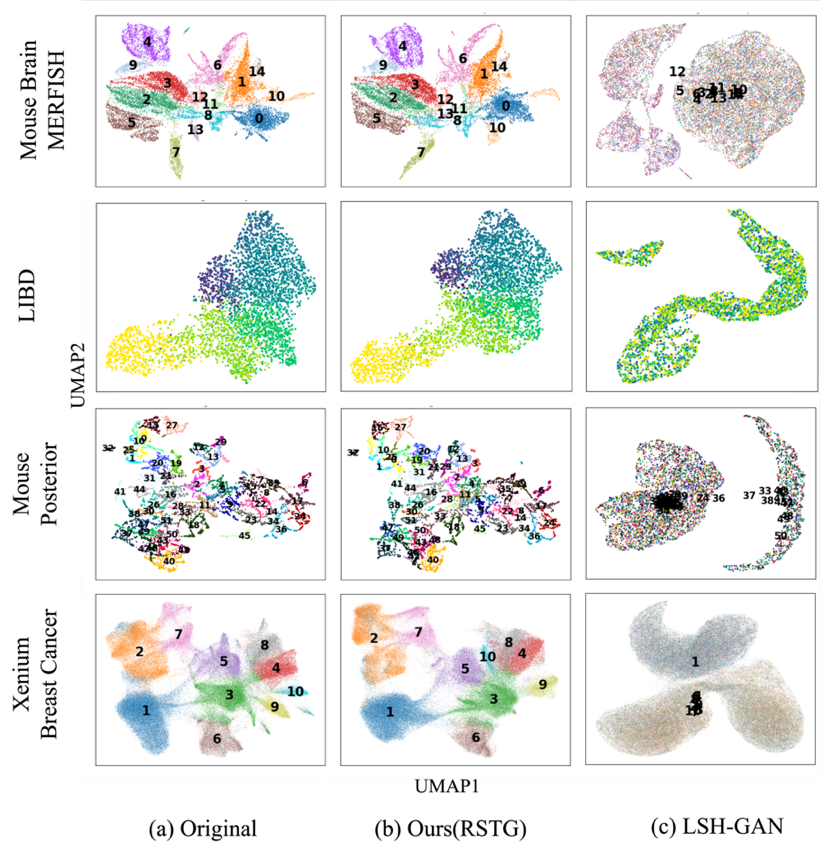

**Supplementary Figure 4.** [a–f] Cortical layer assignment for LIBD tissue IDs 151676 and 151507. [a,d] show true layers; [b,e] show predicted layers. [c,f] present top-1 and top-2 accuracy comparisons across methods.

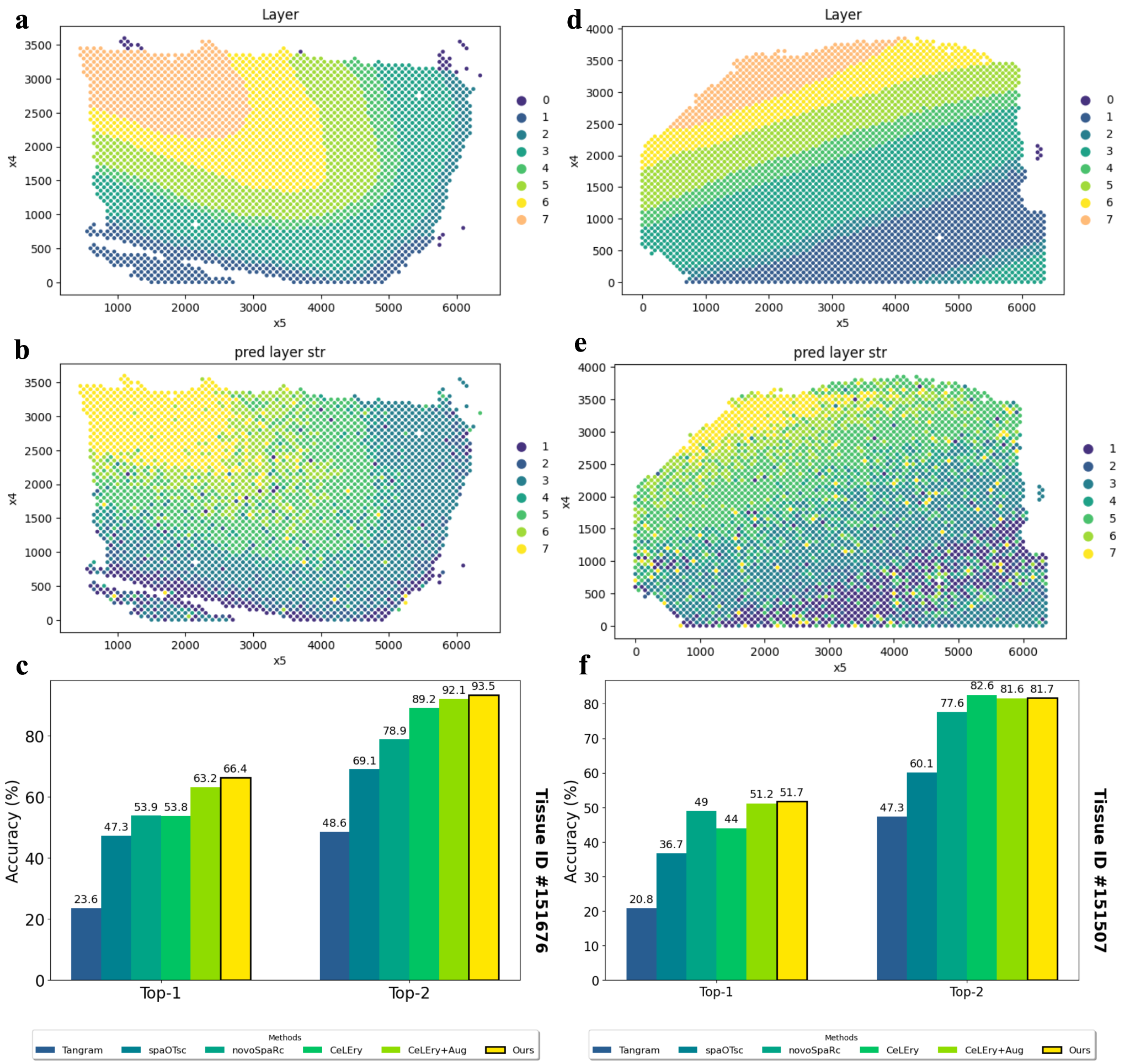

**Supplementary Figure 5.** Gene expression reconstruction and distance correlation on Mouse Posterior dataset. The reconstructed expression map (30% hold-out) illustrates spatial prediction quality. The accompanying bar plot compares Pearson correlation of pairwise distances between predicted and true locations across methods (RSTG, CeLEry, spaotSc, novoSpaRc, Tangram) for beta values 0.005, 0.01, and 0.03, with 10% and 30% test set splits.

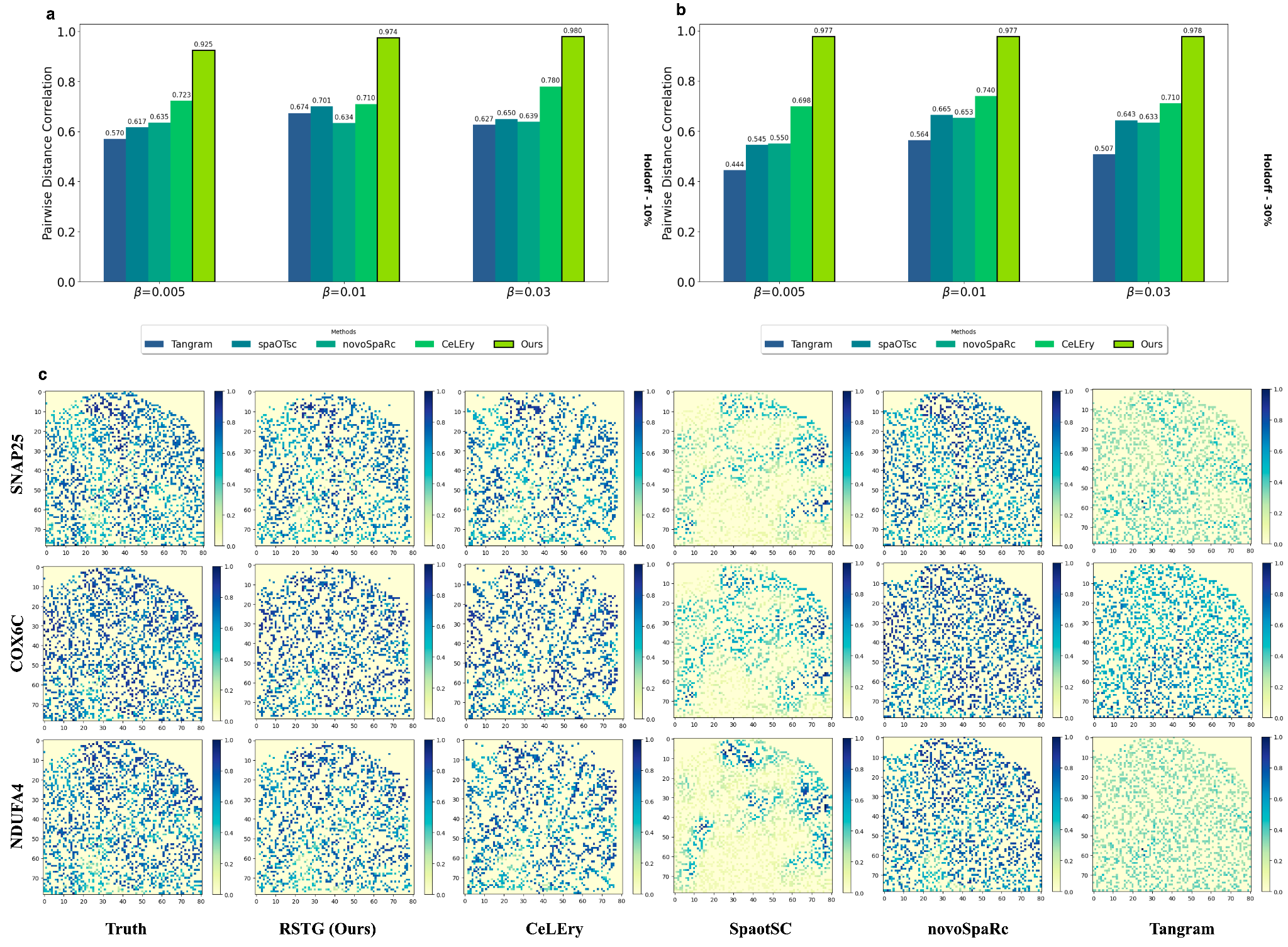

**Supplementary Figure 6.** 2D location recovery on MERFISH mouse brain data. (a) Reconstructed gene expression maps using predicted locations from different methods (Ours, CeLEry, spaotSc, novoSpaRc, Tangram).

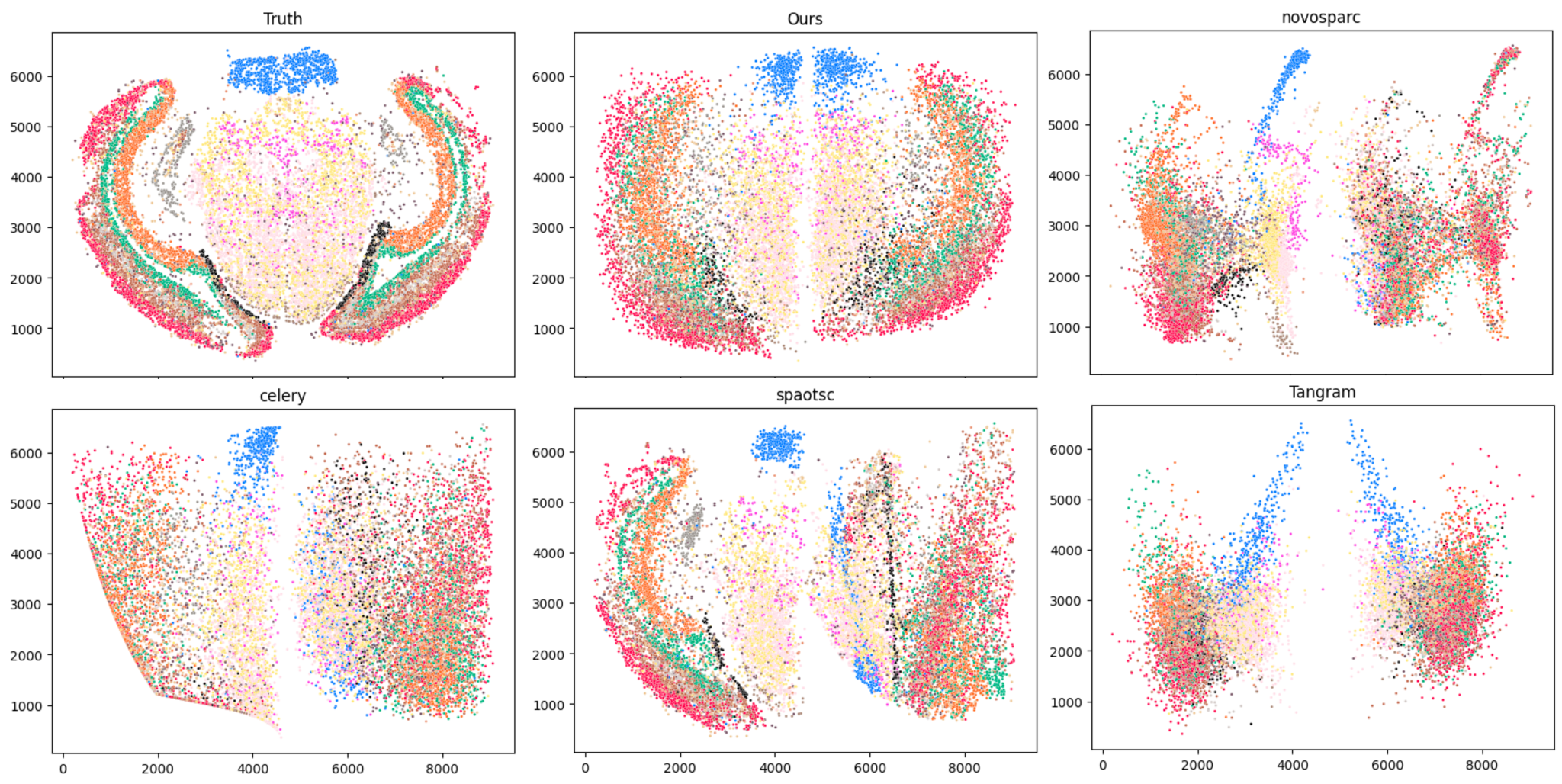
